# A sex difference in the composition of the rodent postsynaptic density

**DOI:** 10.1101/802538

**Authors:** Tara L. Mastro, Anthony Preza, Shinjini Basu, Shona Chattarji, Sally M. Till, Peter Kind, Mary B. Kennedy

## Abstract

SynGAP is a postsynaptic density (PSD) protein that binds to PDZ domains of the scaffold protein PSD-95. We previously reported that heterozygous deletion of synGAP in mice is correlated with increased steady-state levels of other key PSD proteins that bind PSD-95, although the level of PSD-95 remains constant (Walkup et al., 2016). For example, the ratio to PSD-95 of Transmembrane AMPA-Receptor-associated Proteins (TARPs), which mediate binding of AMPA-type glutamate receptors to PSD-95, was increased in young *synGAP*^*+/-*^ mice. Here we show that a highly significant increase in TARP in the PSDs of young *synGAP*^*+/-*^ rodents is present only in females and not in males. The data reveal a sex difference in the adaptation of the PSD scaffold to synGAP heterozygosity.

---

SynGAP is a Ras/Rap GTPase Activating Protein that is specifically expressed in neurons and is highly concentrated in the postsynaptic density (PSD) of glutamatergic synapses in the brain (Chen et al., 1998; Kim et al., 1998). Mutations in the human gene synGAP1 that cause heterozygous deletion or dysfunction of synGAP result in a severe form of intellectual disability (synGAP haploinsufficiency, also called Mental Retardation type 5 [MRD5]) often accompanied by autism and/or seizures (Berryer et al., 2013; Hamdan et al., 2011; Hamdan et al., 2009). In mice, heterozygous deletion of synGAP causes similar neurological deficits; homozygous deletion causes death a few days after birth (Komiyama et al., 2002; Vazquez et al., 2004).

One function of synGAP is to regulate the balance of active Ras and Rap at the postsynaptic membrane (Walkup et al., 2015), thereby controlling the balance of exocytosis and endocytosis of AMPA-type glutamate receptors (Zhu et al., 2002) and contributing to regulation of the actin cytoskeleton (Tolias et al., 2005). In a recent paper in eLife (Walkup et al., 2016), we postulated a second function which involves regulation of anchoring of AMPA-type glutamate receptors (AMPARs) in the PSD. AMPARs are tethered to the scaffold protein PSD-95 in the PSD by auxiliary subunits called TARPs (Transmembrane AMPA Receptor-associated Proteins, Tomita et al., 2003). TARPs contain a PDZ ligand that binds to PDZ domains in PSD-95. An early event in induction of long-term potentiation (LTP) is increased trapping of AMPARs that is mediated by enhanced binding of TARPs to PDZ domains (Tomita et al., 2005). SynGAP is also anchored in the PSD by binding of its α1 splice variant to the PDZ domains of PSD-95 (Kim et al., 1998; McMahon et al., 2012; Walkup et al., 2016). SynGAP is nearly as abundant in the PSD fraction as PSD-95, which suggests that it occupies a large fraction of PDZ domains and can compete with TARPs for binding to PSD-95 (Chen et al., 1998; Dosemeci et al., 2007). During induction of LTP, calcium/calmodulin-dependent protein kinase II (CaMKII) is activated and phosphorylates synGAP, increasing the rate of inactivation of Rap relative to Ras, and, at the same time, causing a decrease in the affinity of synGAP-α1 for the PDZ domains of PSD-95 (Walkup et al., 2015, 2016). We postulated that the decreased affinity of synGAP for PSD-95 might contribute to induction of LTP by allowing TARPs and their associated AMPARs to compete more effectively for binding to the PDZ domains and thus increase their anchoring in the PSD. If this hypothesis is correct, one consequence could be that induction of LTP would be disrupted in synGAP heterozygotes because the transient shift in competition for PDZ binding by synGAP would be less potent as a result of the mutation. A second possible consequence could be that the steady state level of TARPs bound to PSD-95 in PSDs would be increased in synGAP heterozygotes because the steady state level of synGAP is reduced.

In our original study, we tested the second possibility by determining the ratio of TARPs to PSD-95 in PSD fractions prepared from pooled forebrains of *synGAP*^*+/-*^ (HET) and of *wild type* (WT) mice (5 WT males and 1 WT female; 4 HET males and 2 HET females, ranging in age from 7.9 weeks to 12.4 weeks) The ratio of synGAP to PSD-95 was 25% reduced in PSDs from the HET mice compared to WT. As we had predicted, the average ratio of TARPs to PSD-95 showed a small (12%) but significant increase in PSDs from the HET animals compared to WT (Walkup et al., 2016). We also found a small but significant increase in the ratio of LRRTM2 (14%) and neuroligin-2 (9%) to PSD-95. The ratio of neuroligin-1 to PSD-95 was unchanged.

To attempt to reproduce these findings with a larger data set and to examine the biological variability in the relationship between the amount of synGAP and the amount of TARP in PSD fractions, we devised a method to isolate PSD fractions from individual WT and HET rodents and we measured the ratios of synGAP and TARPs to PSD-95 in each individual PSD fraction. We also measured the ratios of GluN2B, neuroligin1, and neuroligin2 to PSD-95. We were able to study mice and rat HETs by using a new rat mutant in which one copy of the synGAP gene is inactivated by the CRISPR-Cas9 method.

When the data was averaged over all WT and HET animals in this large data set, the TARP/PSD-95 ratio in PSDs was not different between WT and HET animals. Nonetheless, we confirmed our earlier conclusion by using the more powerful Spearman’s rank correlation coefficient to show a statistically significant inverse correlation between the synGAP/PSD-95 and TARPs/PSD-95 ratios in individual PSD fractions from HET rodents. However, to our surprise, when data from HET rodents are separated by sex, the inverse correlation is present only in females and not in males. The large and highly significant inverse correlation in HET females drives a significant inverse correlation in the pools of all HET animals and of all female animals. The inverse correlation is not found in any subset of animals that contains only males.

## Results

### Creation of rat synGAP KO by the CRISPR-Cas9 method

CRISPR/Cas9 technology was used to establish a *SynGAP1* KO rat line that harbors a frameshift mutation in exon8 of *SynGAP1* (Fig. 1A), which prevents expression of the protein. SynGAP protein expression level is reduced by 50% in HET knockout rats compared to wild-type (WT) and is absent in homozygous knockouts (Fig. 1B). While SynGAP KO rats die perinatally, SynGAP HET rats appear healthy and fertile.

**Fig. 1.**
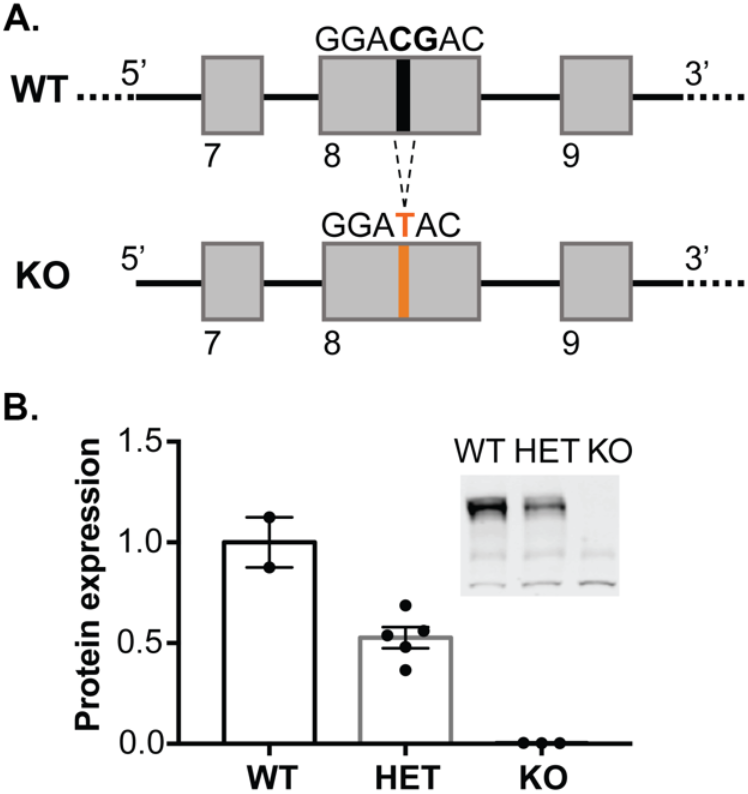
Generation of synGAP null rats. A) SynGAP targeting strategy introduces a frame shift mutation into exon 8. B) Quantification of synGAP immunoblots (inset) of cortical homogenates was performed as described in Methods.

### Average synGAP/PSD-95 and TARP/PSD-95 ratios in WT and HET rodents

PSD fractions were prepared from the forebrains of 165 individual rodents, comprising 82 WT and 83 HETs, as described under Materials and Methods. The total sample included 81 females (39 WT, 42 HET), and 84 males (43 WT, 41 HET). In each category, approximately half of the animals were rats and half were mice; approximately half were 7.5 weeks old and half were 12.5 weeks old. The ratio of synGAP/PSD-95 and TARP/PSD-95, averaged over all of the rodents, are summarized in the two bars labeled “All” (Fig 2A,B, left). As expected, the synGAP/PSD-95 ratio (Fig. 2A, left) is reduced by 22% in HET rodents compared to WT (the WT level is indicated by a dotted line). However, the ratio of TARP to PSD-95 (Fig. 2B, left) is not significantly changed. The results were similar for averages of animals grouped by sex, species, and age (Fig. 2 A and B), except for 7 wk old female mice in which the ratio of TARPs to PSD-95 was significantly reduced compared to WT. This value may have been influenced by lower overall expression of TARPS in 7 week old mice. We also noted more variability in the averaged ratios of TARP to PSD-95 for females (Fig 2B, right) compared to males (Fig. 2B, middle). Taken as a whole, it appears that the averaged results do not reproduce our original finding in Walkup et al. (2016).

**Figure 2.**
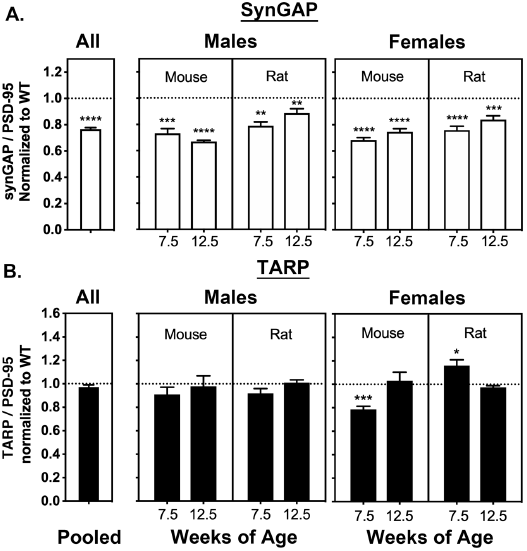
Averaged ratios of synGAP and TARPs to PSD-95 in PSDs from WT and HET Rats and Mice. PSDs were purified from the brains of individual animals as described under Methods. The ratios of synGAP to PSD-95 (A) and TARPs to PSD-95 (B) were determined as described under Methods. Ratios from HET animals (bars) are normalized to the ratios from WT animals (dotted lines). Antibodies against synGAP, TARPS, and PSD-95 are the same as those used in (Walkup et al., 2016). The antibody against synGAP (AB_2287112) recognizes all isoforms of synGAP. The antibody against TARPs (AB_877307) recognizes TARP-γ2, γ3, γ4, and γ8. The sample sizes for each group and the significance tests are as follows: A) all animals WT=79 and HET=78, one-tailed Wilcoxon matched-pairs signed rank test; male mouse 7.5 weeks WT=11 and HET=9, one-tailed Student T-test; male mouse 12.5 weeks WT=11 and HET=8, one-tailed Student T-test with Welch’s correction; male rat 7.5 weeks WT=11 and HET=10, one-tailed Student T-test; male rat 12.5 weeks WT=10 and HET=11, one-tailed Student T-test; female mouse 7.5 weeks WT=10 and HET=12, one-tailed Student T-test with Welch’s correction; female mouse 12.5 WT=9 and HET=9, one-tailed Student T-test; female rat 7.5 weeks WT=10 and HET=10, one-tailed Student T-test; female rat 12.5 weeks WT=9 and HET=10, one-tailed Student T-test. B) all animals WT=77 and HET=80, one-tailed Wilcoxon matched-pairs signed rank test; male mouse 7.5 weeks WT=10 and HET=9, one-tailed Student T-test; male mouse 12.5 weeks WT=10 and HET=10, one-tailed Mann Whitney test; male rat 7.5 weeks WT=10 and HET=10, one-tailed Student T-test; male rat 12.5 weeks WT=10 and HET=1, one-tailed Student T-test; female mouse 7.5 weeks WT=9 and HET=10, one-tailed Student T-test; female mouse 12.5 WT=9 and HET=10, one-tailed Mann Whitney test; female rat 7.5 weeks WT=10 and HET=10, one-tailed Student T-test; female rat 12.5 weeks WT=9 and HET=10, one-tailed Student T-test with Welch’s correction. Significance: * for p ≤ 0.05, ** for p ≤ 0.01, *** for p ≤ 0.001, and **** for p ≤ 0.0001.

### Spearman’s correlation coefficient reveals that the synGAP/PSD-95 and TARP/PSD-95 ratios are inversely correlated only in females

Comparison of averaged ratios is not a perfect measure of the correlation between two ratios among individuals. Therefore, we used a more direct measure, Spearman’s rank correlation coefficient r, to test for correlation between the synGAP/PSD-95 and TARP/PSD-95 ratios among individuals. This statistic measures whether a monotonic correlation exists between the rank orders of the magnitudes of two variables in a data set. If the ranks of the two variables correlate perfectly, Spearman’s r is 1; if there is no correlation, it is zero; and if the ranks are perfectly anti-correlated, it is -1.

We first established that the amounts of PSD-95 per PSD protein are not statistically different between HETs and WT or between male and female subgroups (Fig. 3S1). Thus, differences among individuals in the target protein/PSD-95 ratio can be interpreted as differences in the concentrations of the target protein in the PSD fractions.

A comparison of rank orders among individuals across the four cohorts (7.5 week old mice, 7.5 week old rats, 12.5 week old mice, and 12.5 week old rats) required an additional normalization. The average intensities of staining for proteins differed significantly between the cohorts, presumably because of developmental changes in protein expression. We therefore normalized the ratios for all cohorts to account for these average differences, as described under Methods. The normalization enabled us to look for correlations between ratios among individuals across cohorts.

Fig. 3 contains scatter plots of the TARP/PSD-95 ratio against the synGAP/PSD-95 ratio in individual PSDs from the indicated data sets. Fig. 3A contains the plot for all 165 animals. Rows two (B., E., and H.) and three (C., F., and I.) show separate plots for WT and HET animals, respectively; columns two (D., E., and F.) and three (G., H., and I) show separate plots for females and males, and data from HET animals are indicated in orange to highlight that they have a lower average synGAP/PSD-95 ratio than WT animals. In each plot, Spearman’s r is given, along with the p-value indicating the probability that Spearman’s r differs from zero. P-values that indicate statistical significance are in red. For reference, red lines through the points in each graph show the best linear fit determined by regression. There is not a statistically significant inverse correlation (negative Spearman’s r) between the two ratios for the group of all animals (Fig. 3A).

**Figure 3.**
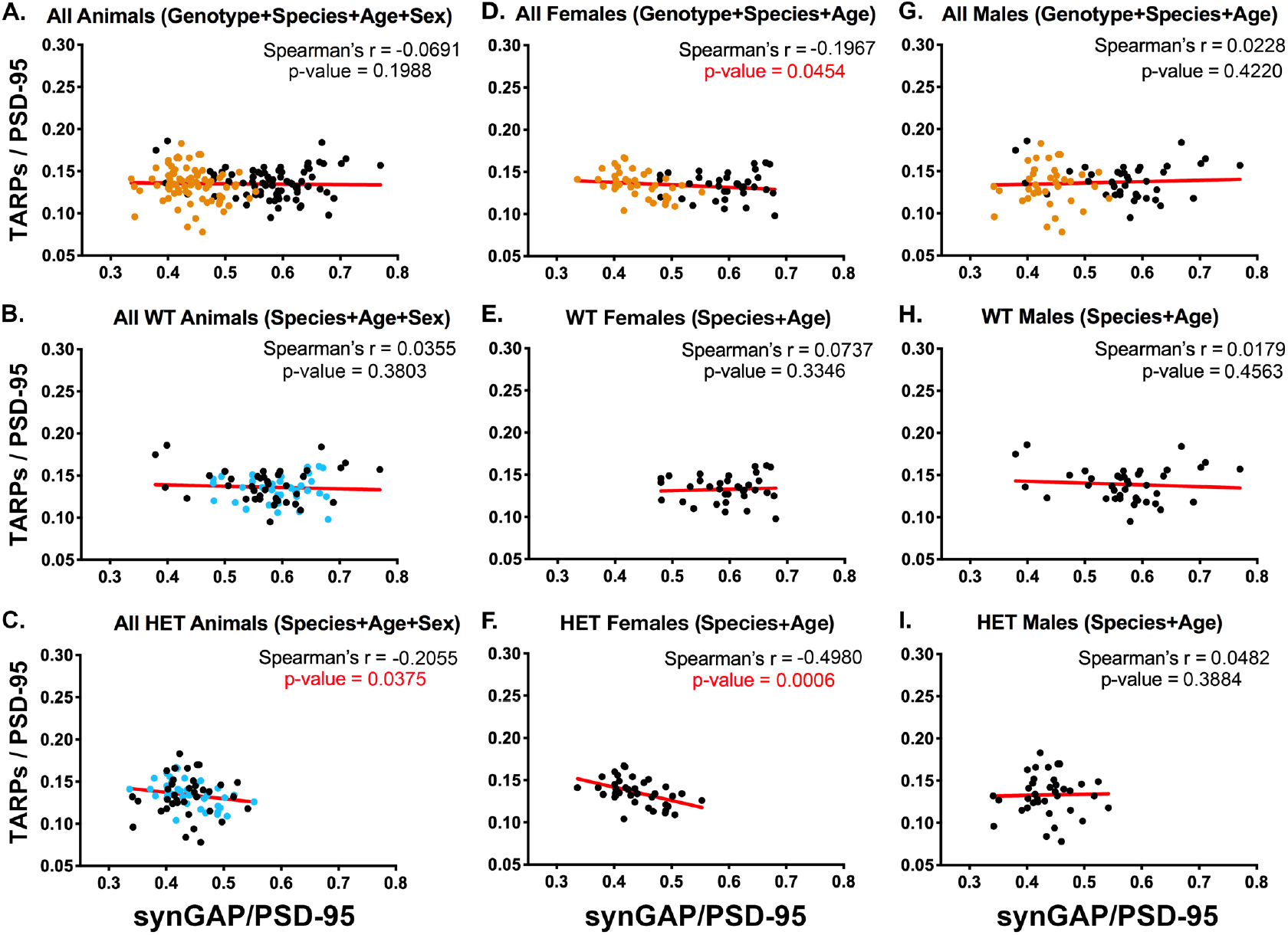
Correlation of the ratios TARPs/PSD-95 and synGAP/PSD-95 among individual animals. Each point represents a single animal. Corrected ratios and Spearman’s rank correlation coefficients were determined as described under Methods. A) All animals, including all genotypes, ages, species, and sexes. Black, WT; Orange, HET; n = 152. B) All WT animals, including all ages, species, and sexes. Black, males; Blue, females. n = 76. C) All HET animals, including all ages, species, and sexes. Black, males; Blue, females; n = 76. D) All female animals, including all genotypes, ages, and species. Black, WT; Orange, HET; n = 75. E) All WT females including all ages and species. n = 36. F) All HET females, including all ages and species. n= 39. G) All male animals, including all genotypes, ages, and species. Black, WT; Orange, HET. n = 77. H) All WT males, including all ages and species. n = 40. I) All HET males, including all ages and species. n = 37. Significant p-values for Spearman’s rank correlation coefficient are shown in red.

Figs. 3B, and C contain plots for all WT and all HET animals, respectively, with ratios from female animals indicated in blue. These data show that, at steady state *in vivo*, lower amounts of synGAP in PSDs from the HET animals (Fig. 3C) correlate with higher amounts of TARP; whereas there is no correlation in WT animals (Fig. 3B). This finding supports our original report (Walkup et al., 2016).

The data from all females (Fig. 3D) show an inverse correlation between the two ratios that just reaches statistical significance. In contrast, the data from all males (fig. 3G) shows no correlation. Similarly, WT females and WT males (Figs. 3E and H) show no correlation. Strikingly, the data from HET females (Fig. 3F) show the largest inverse correlation of all the data sets, with a Spearman’s r = - 0.498 and a p-value = 0.0006. HET males (Fig. 3I) show no significant correlation. The strong inverse correlation between the amount of synGAP and the amount of TARP in PSDs from HET females (Fig. 3F) likely drives the inverse correlation observed for pooled HET animals (Fig. 3C) and pooled females (Fig. 3D). These results mean that, between 7.5 and 12.5 weeks of age, synGAP heterozygosity has a much greater effect on the content of TARPs in the PSDs of female animals than in those of males. The simplest explanation for this is that in HET females, the structure of the PSD, which is determined by multiple equilibria among several proteins, is such that TARP and synGAP compete directly for bindingto PSD-95; whereas in HET males, this particular competition is not significant. Possible underlying mechanisms are outlined in the Discussion.

We also compared data sets from mice and rats at 7.5 weeks and 12.5 weeks (Fig. 3S2). These data sets were small (9 or 10 animals). Nevertheless, they show a statistically significant inverse correlation between TARP/PSD-95 and synGAP/PSD-95 in HET female mice at both 7.5 and 12.5 weeks. In data from HET rats at 7.5 weeks, the inverse correlation is very close to significance; at 12.5 weeks, it is less significant, but still shows a trend. In the corresponding males, none of the data sets shows a statistically significant inverse correlation. More data would be required to make a definitive conclusion, but the results suggest that competition between synGAP and TARP for binding to PSD-95 in females is more prominent at 7 weeks than at 12 weeks, and more prominent in mice than in rats.

### Effect of synGAP haploinsufficiency on the relative levels of other PSD proteins

In our previous paper, we examined the levels of neuroligins 1 and 2 (NLG-1, -2), and of the surface protein LRRTM2. In this study, we re-examined the effect of reduction of synGAP on the levels of NLG-1 and 2 in the PSD and looked at the effect on levels of GluN2B, a subunit of the NMDA-type glutamate receptor that binds most avidly to PDZ2 of PSD-95. We predicted that the level of GluN2B would be less affected than TARPs or NLGs by reduction of synGAP because synGAP has lower affinity for PDZ2 than for PDZ1 and PDZ3 (Walkup et al., 2016).

The ratios of the three proteins to PSD-95 in HET and WT rodents, averaged over the same PSD fractions shown in Figs. 2A and B, are shown in the bars labeled “All” in Fig. 4A, B, and C (left). GluN2B exhibits a highly significant reduction of about 10% in HETs compared to WT; NLG-1 shows no change; and NLG-2 increases significantly by about 7%. There is no significant difference in these ratios between rat and mouse, males and females, or between 7.5 week and 12.5 week old animals. The absence of any change in NLG-1 and the slight increase in NLG-2 recapitulate our findings in Walkup et al. (2016).

**Fig. 4.**
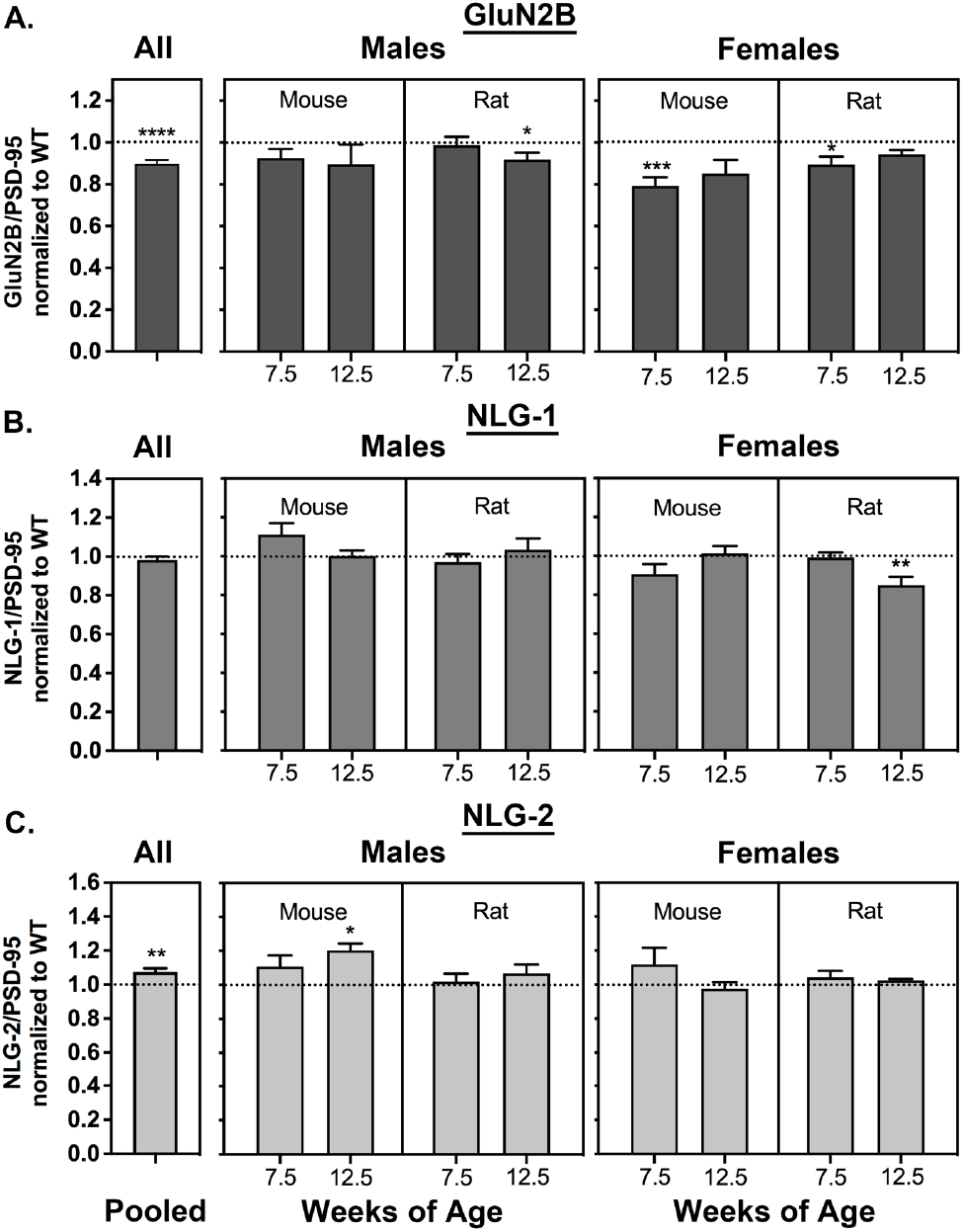
Averaged ratios of GluN2B, NLG1, and NLG-2 to PSD-95 in synGAP mice and rat HETs. PSDs were purified as in Fig. 2. Ratios were determined as described under Methods. Ratios from HET animals (bars) are normalized to the ratios from WT animals (dotted lines). A) GluN2B/PSD-95. Sample sizes and significance tests are as follows: all animals WT=81 and HET=82, one-tailed Wilcoxon matched-pairs signed rank test; male mouse 7.5 weeks WT=11 and HET=10, one-tailed Student T-test; male mouse 12.5 weeks WT=11 and HET=10, one-tailed Student T-test; male rat 7.5 weeks WT=11 and HET=10, one-tailed Student T-test; male rat 12.5 weeks WT=10 and HET=11, one-tailed Student T-test; female mouse 7.5 weeks WT=10 and HET=12, one-tailed Student T-test; female mouse 12.5 WT=9 and HET=9, one-tailed Student T-test; female rat 7.5 weeks WT=9 and HET=10, one-tailed Student T-test with Welch’s correction; female rat 12.5 weeks WT=9 and HET=9, one-tailed Student T-test with Welch’s correction. B) NLG-1/PSD-95. Sample sizes and significance tests are as follows: all animals WT=81 and HET=83, one-tailed Wilcoxon matched-pairs signed rank test; male mouse 7.5 weeks WT=10 and HET=10, one-tailed Student T-test; male mouse 12.5 weeks WT=11 and HET=10, one-tailed Student T-test; male rat 7.5 weeks WT=11 and HET=10, one-tailed Student T-test; male rat 12.5 weeks WT=10 and HET=11, one-tailed Student T-test; female mouse 7.5 weeks WT=10 and HET=12, one-tailed Student T-test; female mouse 12.5 WT=10 and HET=10, one-tailed Student T-test; female rat 7.5 weeks WT=10 and HET=11, one-tailed Mann-Whitney test; female rat 12.5 weeks WT=9 and HET=10, one-tailed Student T-test with Welch’s correction. C) NLG-2/PSD-95. Sample sizes and significance tests are as follows: all animals WT=79 and HET=79, one-tailed Wilcoxon matched-pairs signed rank test; male mouse 7.5 weeks WT=10 and HET=10, one-tailed Student T-test; male mouse 12.5 weeks WT=11 and HET=10, one-tailed Student T-test; male rat 7.5 weeks WT=11 and HET=10, one-tailed Student T-test; male rat 12.5 weeks WT=10 and HET=11, one-tailed Student T-test; female mouse 7.5 weeks WT=9 and HET=12, one-tailed Student T-test with Welch’s correction; female mouse 12.5 WT=10 and HET=9, one-tailed Student T-test; female rat 7.5 weeks WT=9 and HET=10, one-tailed Student T-test; female rat 12.5 weeks WT=9 and HET=8, one-tailed Student T-test with Welch’s correction. Significance: * for p ≤ 0.05, ** for p ≤ 0.01, *** for p ≤ 0.001, and **** for p ≤ 0.0001.

### Spearman’s correlation coefficient reveals that the levels of GluN2B and levels of synGAP in PSDs are positively correlated

Fig. 5 contains ratios of GluN2B to PSD-95 plotted against ratios of synGAP to PSD-95 measured in the same set of individual PSDs shown in Fig 3. A, D, and G. Data for all animals (Fig. 5A), female animals (Fig. 5B), and male animals (Fig. 5C) show positive Spearman’s r values of ∼ 0.4 with p-values indicating a highly significant difference from zero. The positive correlation is present in both WT (black) and HET (orange) animals. These results support our hypothesis that synGAP does not compete with GluN2B for binding to PSD-95. To the contrary, the data suggest that a higher level of synGAP leads to higher localization of GluN2B; and, therefore, NMDA-type glutamate receptors, to the PSD (see Discussion).

**Figure 5.**
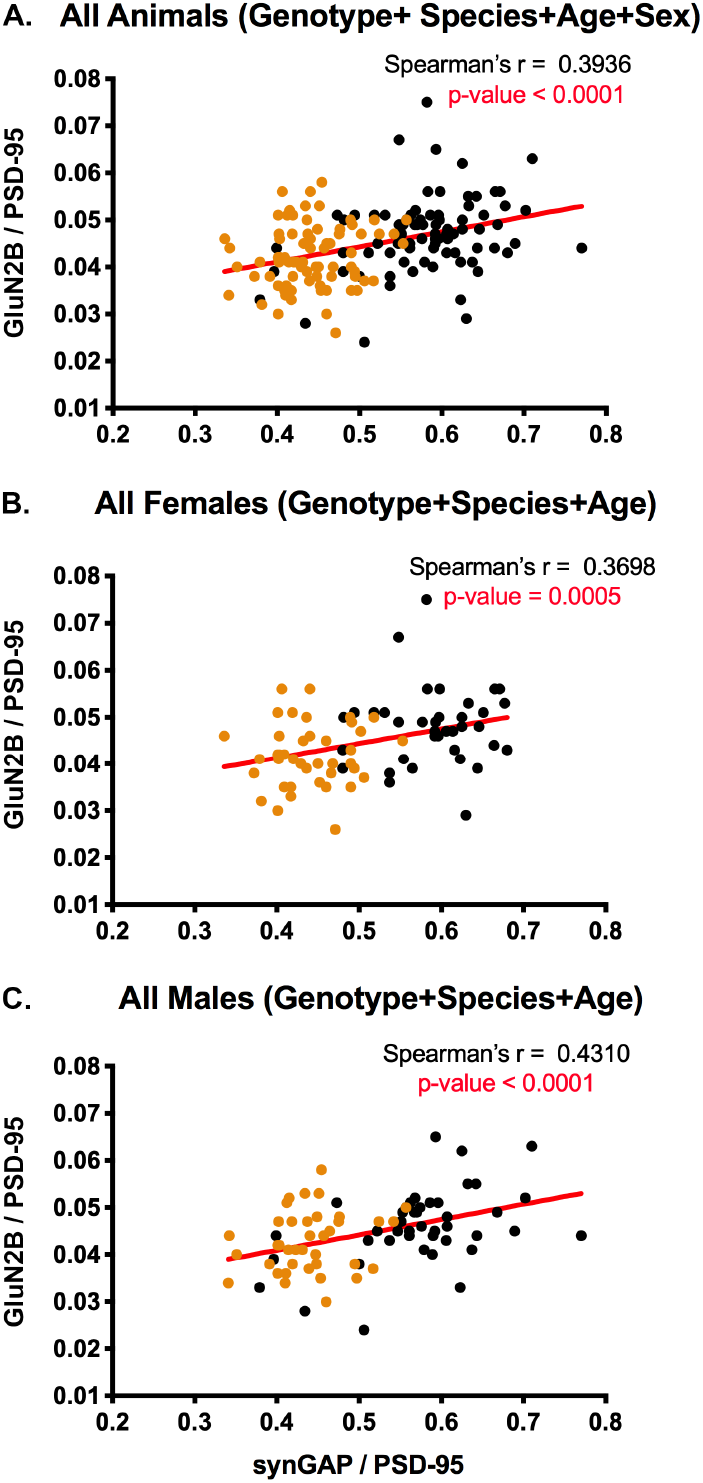
Correlation of the ratios GluN2B/PSD-95 and synGAP/PSD-95 for individual animals. Each point represents a single animal. Black, WT; Orange, HET. A) All animals including all genotypes, ages, species, and sexes. n = 158. B) All female animals, including all genotypes, ages, and species. n = 77. C) All male animals, including all genotypes, ages, and species. n = 81. Significant p-values for Spearman’s rank correlation coefficient are shown in red.

### Spearman’s correlation coefficient shows no significant correlation between levels of NLG-1 and 2 and levels of synGAP in PSDs

Our previous results showed no effect of synGAP haploinsufficiency on the amount of NLG-1 in PSDs and only a small effect on the level of NLG-2. The pooled data in Fig. 4C and Spearman’s r in Fig. 6B reproduce those findings. However, the significance of Spearman’s r between levels of NLG-2 and synGAP in PSDs shows only a strong trend toward an inverse correlation (Fig. 6A).

**Figure 6.**
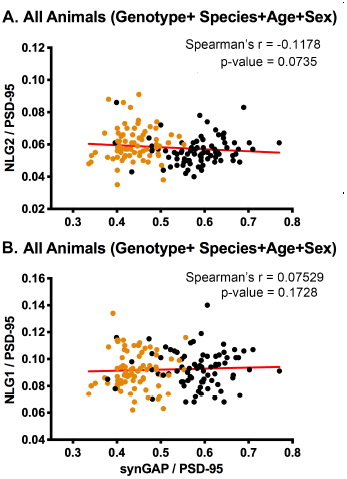
Correlation of the ratios of NLG-2/PSD-95 or NLG-1/PSD-95 and synGAP/PSD-95 for individual animals. Each point represents a single animal. Black, WT; Orange, HET. A) Correlation of NLG-2/PSD-95. All animals, including all genotypes, ages, species, and sexes. n = 153. B) Correlation of NLG-1/PSD-95. All animals, including all genotypes, ages, species, and sexes. n = 159. P-values of Spearman’s rank correlation coefficient indicate no significant correlation.

## Discussion

The most striking new result of this study is the discovery of a sex difference in the adaptation of the PSD scaffold to synGAP haploinsufficiency. Specifically, we show that a decrease in the steady-state concentration of synGAP in rodent PSDs correlates with a higher concentration of TARPs in PSDs only in females and not in males. In female HETs, the rank correlation coefficient between the concentrations of TARP and synGAP in PSDs is -0.5, which suggests a relatively high competition between the two proteins for binding to PDZ domains of PSD-95 *in vivo*. This competition does not affect the composition of PSDs in male HETs.

SynGAP haploinsufficiency causes about 4% of cases of sporadic intellectual disability (ID) in humans, often accompanied by seizures and autistic behaviors (Berryer et al., 2013). If the sex difference in the interaction of synGAP with TARPs in the rodent PSD is also present in humans, it might result in differences between girls and boys in the prevalence of some of the associated symptoms. The PSD is formed by multiple interactions among the major scaffold proteins and “client” signaling proteins which are concentrated in the PSD by their association with the scaffold proteins (Kennedy, 2013; Kennedy et al., 2005; Sheng and Kim, 2011).

At any one time, the composition of a PSD is a dynamic equilibrium among all the possible protein associations, driven by the relative concentrations of each protein in a spine, and the relative affinities of their mutual binding domains (e.g. Gray et al., 2006). The simplest interpretation of our present result is that there is a difference between males and females in the composition or regulation of proteins in PSDs which causes the concentration of TARPs in PSDs to be sensitive to the steady-state concentration of synGAP in females, but not males. This occurs despite the fact that there is no difference between males and females in the amount of synGAP in PSDs of either genotype.

In our recent eLife paper, we postulated that synGAP-α1, which contains a PDZ ligand, competes with TARPs for binding to PSD-95 and therefore helps to limit the number of AMPARs immobilized at the synapse (Walkup et al., 2016). This hypothesis has two possible corollaries. One is that transient phosphorylation of synGAP by CaMKII during induction of LTP, which reduces the affinity of its PDZ ligand for PSD-95, will allow more binding of TARP to the PDZ domains; and thus contribute to increased trapping of AMPARs (see Opazo and Choquet, 2011; Tomita et al., 2005). The second corollary, which is addressed in this study, concerns the steady-state composition of PSDs in synGAP^+/-^ rodents. If, at steady-state, the concentration of synGAP in WTs is high enough to effectively compete with binding of TARPs to PSD-95, then, in HET rodents, which have a reduced concentration of synGAP, the concentration of TARPs (and thus AMPARs) in the PSD will be higher than in WT. Indeed, we previously reported a higher concentration of TARPs in PSDs isolated from a pool of six HET mice compared to a pool of six WT mice. The pools were approximately matched for age and sex; but the HET pool contained two females, whereas the WT pool contained one (Walkup et al., 2016). Because the increase in concentration of TARPs in the HET pool was significant but small, we decided to examine the correlation of TARP and synGAP concentrations in a large set of individual PSDs. To do this, we developed a method for isolating PSDs from individual rodents. The analysis supports the hypothesis that synGAP-α1 does indeed compete with steady-state binding of TARPs to PSD-95 in some circumstances.

There are many possible mechanistic explanations for the sex difference in sensitivity of TARPs to the concentration of synGAP. One simple one is that additional protein(s) are present in females that compete with synGAP for binding to PDZ domains of PSD-95. The resulting “crowding” could make binding of TARP to PSD-95 more sensitive to loss of synGAP in females.

Another is that an additional protein which can compete with TARP more strongly than synGAP for binding to PSD-95 is present in male PSDs, but not in female PSDs. In this case, reduction of synGAP in PSDs of males would be expected to have little effect on the concentration of TARPs. Interestingly, despite any differences between males and females in mechanism, the mean steady-state ratio of TARPs to PSD-95 in the two sexes is not significantly different among WTs or HETs.

Two recent studies have documented sex differences in regulation of synaptic plasticity. Wang et al. (2018) found that 7 to 12 week old female rodents (the same age range used in our study) have higher synaptic levels of membrane estrogen receptor alpha (mERalpha) and, in contrast to males, require activation of mERalpha for activation of some of the signaling kinases that support long-term potentiation. This difference results in a higher threshold in females for LTP and for some forms of spatial learning. Jain et al. (2019), studying hippocampal slices from 7 to 10 week old rats, found that estrogen-induced plasticity, which occurs in both males and females, requires synergistic activation of L-type Ca^2+^ channels and internal Ca^2+^ stores in females; whereas in males, either of the two sources is sufficient. In addition, activity-dependent LTP requires activation of protein kinase A in females, but not in males. They conclude that there are latent sex differences in mechanisms of synaptic potentiation in which distinct molecular signaling pathways converge to common functional endpoints in males and females. Neither of these studies provides an immediate explanation for our findings, but they support the idea that there are a number of biochemical and structural differences between males and females in the synaptic regulatory apparatus of 7 to 12 week old rodents.

Because the cytosolic tail of the GluN2B subunit binds to the first and second PDZ domains of PSD-95 (Kornau et al., 1995), we tested whether the concentration of GluN2B in PSDs is altered by synGAP haploinsufficiency. In contrast to TARPs, the concentration of GluN2B in PSDs shows a strong positive correlation with the amount of synGAP (rank correlation coefficient ≈ 0.4), and is reduced in synGAP HETs. The rank correlation among individual animals is significant in both WT and HET animals and does not differ between males and females (Fig. 5). This data shows that synGAP plays a role in localizing GluN2B to the PSD, but that it is not the only protein involved.

We reproduced our original finding that the amount of synGAP in the PSD does not influence the amount of NLG-1 and shows a trend toward a small inverse correlation with the amount of NLG-2 in PSDs (Walkup et al., 2016). Thus, although both synGAP and NLG-1 (Irie et al., 1997) bind to PDZ3 of PSD-95, reduction of synGAP does not increase the steady-state amount of NLG-1 in the PSD *in vivo*. The simplest explanation is that NLG’s have a higher affinity for PDZ3 than synGAP such that synGAP does not compete effectively with them for binding to PDZ3 *in vivo*. The affinities of NLG-1 and of synGAP for PSD-95 have been estimated by Biacore surface plasmon resonance technology; but, the measurement parameters were not directly comparable. A protein fragment containing all three PDZ domains of PSD-95 was found to have a K_D_ of ∼ 200 nM for a 16mer with the sequence of the carboxyl terminus of NLG-1 (Irie et al., 1997). In our recent study, we found that nearly full length synGAP, missing only the first 100 residues of the amino-terminus, has an affinity for PDZ3 of ∼650 nM and for a fragment containing all three PDZ domains of 5 nM. It is likely that the competition for binding to PDZ3 between NLG’s and synGAP is influenced by additional binding sites on PSD-95 or on other scaffold proteins for NLG’s and/or synGAP.

These results illustrate the complex role that synGAP plays in modulating the steady-state composition of the PSD. It can compete with some proteins for localization in the PSD and it can help to concentrate others. The results are consistent with the concept that the structure of the PSD is a dynamic equilibrium governed by multiple protein associations and driven by the relative concentrations of each protein and the affinities of their mutual binding domains. This study has not tested the effect of the concentration of synGAP on the acute re-organization of the PSD that occurs, for example, upon induction of LTP. Our earlier finding that phosphorylation of synGAP by CaMKII, which is activated upon induction of LTP, reduces the affinity of synGAP for all three of the PDZ domains of PSD-95 suggests that post-translational modulation of synGAP’s affinity for PSD-95 may initiate transient reorganization of the PSD in stimulated synapses. Disruption of this transient role likely contributes to the phenotypes of synGAP haploinsufficiency, but in ways that may not be different between females and males.

## Methods

### Animals

SynGAP KO mice were generated in the Kennedy lab, bred in the Caltech animal facility, and genotyped by polymerase chain reaction as described (Vazquez et al., 2004). The LE-*SynGAP1*^*em1/PWC*^, hereafter referred to as *SynGAP* KO, rat model was produced by SAGE Labs, Sigma-Aldrich (now a subsidiary of Horizon Discovery, Saint Louis, MO, USA) using CRISPR/Cas9 based genome targeting strategies (Li et al., 2013). Briefly, CRISPR/Cas9 reagents and sgRNA targeting exon 8 (gtgcatagagcatgtcgtccAGG) were microinjected into pronuclei of fertilized one-cell embryos from Long Evans rats and transplanted into pseudo-pregnant females. Live born pups were genotyped by Sanger sequencing of exon8 PCR amplicons. Founder #23 displayed a 2bp deletion 1bp insertion and was crossed to a wild type Long Evans rat to generate a cohort of heterozygous animals for breeding purposes. Tissue biopsies from pups were used for TaqMan genotyping using two primers (Forward: CCAAGAAGCGATATTACTGCGAGTT and Reverse: GGAAGTGGTCCGTGCATAGA) and two reporter probes (WT: CCTGGACGACATGC and KO: TGCCTGGATACATGC). Homozygous *SynGAP* KO rats die perinatally, but heterozygous (HET) SynGAP KO rats appear healthy and are fertile. To verify knockout of expression of synGAP, forebrain homogenates of P5 littermate rats (n KO=3, n HET=5, n WT=2) were prepared in radioimmunoprecipitation assay buffer containing protease and phosphatase inhibitors (cOmplete EDTA-free), immunoblotted with a primary antibody raised to panSynGAP (1:4000; Abcam AB77235), and imaged on an Odyssey infrared imaging system (Li-COR Bioscience) as previously described in (Till et al., 2012). Total protein levels on the blot were visualised with the Pierce Reversible Protein Stain Kit (Thermo Fisher Scientific, #24580) and quantified with ImageJ gel analyzer software. Protein bands of interest were quantified with Image Studio Lite v5.0 (Li-COR Bioscience). The expression level of each protein of interest was first normalized to total protein, followed by normalization of the data to the average wild-type levels, which were considered to be 100%.

Upon weaning each animal was given an ear punch ID associated with a unique ID number, sexed, and genotyped. Tissue from the ear punch was used for genotyping both mice and rats, although some mice were genotyped with tissue from the tail. Mice were genotyped by polymerase chain reaction as described (Vazquez et al., 2004) and rats were genotyped via sequencing performed by Transnetyx, Inc. Cordova, TN. The ID numbers linked the genotype and all other metadata for an animal to its PSD samples.

### Preparation of PSD Fractions

PSD fractions were prepared from individual WT and HET mice and rats that were either 7.5 weeks or 12.5 weeks old, by a modification of a standard method (Cho et al., 1992; Cohen et al., 1977). The ID numbers were used to label, and track tissue samples and extracts after harvesting from the animal. Animals were killed by decapitation according to a protocol approved by the Caltech Institutional Animal Care and Use Committee. The following steps were carried out at 4° C. Forebrains were dissected from each animal, rinsed in Buffer A (0.32 M sucrose, 1 mM NaHCO_3_, 1 mM MgCl_2_, 0.5 mM CaCl_2_, 0.1 mM phenylmethylsulfonyl chloride [PMSF, Sigma Millipore, St. Louis, MO]). Each individual forebrain was homogenized in Buffer A (10% w/v, 4.5 ml for mice and 13.5 ml for rats) with 12 up and down strokes of a teflon/glass homogenizer at 900 rpm. Homogenates were subjected to centrifugation at 1400 × g for 10 min. The pellet was resuspended in Buffer A to 10% w/v (3.8 ml for mice, 12 ml for rats), homogenized (three strokes at 900 rpm) and subjected to centrifugation at 710 g for 10 min. The two supernatants were combined and subjected to centrifugation at 13,800 g for 10 min. The resulting pellet was resuspended in Buffer B (0.32 M sucrose, 1 mM NaHCO_3_; 2ml for mice, 8 ml for rats), homogenized with 6 strokes at 900 rpm in a teflon/glass homogenizer, and layered onto a discontinuous sucrose gradient (equal parts 0.85 M, 1.0 M, and 1.2 M sucrose in 1 mM NaH_2_CO_3_ buffer; 10.5 ml for mice, 30 ml for rats). Gradients were subjected to centrifugation for 2 hours at 82,500 g in a swinging bucket rotor. The synaptosome-enriched layer at the interface of 1.0 and 1.2 M sucrose was collected, diluted with Buffer B, 7 ml for mice and 20 ml for rats, then added to an equal volume of Buffer B containing 1% Triton (10% X-100 Surfact-Amps, Thermo Fisher Scientific, Waltham, MA). The mixture was stirred for 15 min at 4°C and subjected to centrifugation for 45 min at 36,800 g. The pellet, which contained the PSD-enriched Triton-insoluble fraction, was resuspended in 0.5 -1.0 ml 40 mM Tris pH 8 with a 21-gauge needle attached to a 1 ml syringe, and then homogenized with six strokes at 900 rpm in a teflon/glass homogenizer. Samples were flash-frozen, and stored at -80° C. Protein concentrations were determined by the bicinconic acid method (Thermo Fisher Scientific). Individual mouse brains yielded ∼1.25 mgs and individual rat brains yielded ∼3.5 mgs protein. The yield was sufficient to measure the ratio of amounts of five separate proteins to PSD-95 for each individual animal.

### Immunoblots

To measure each of the five proteins, an equal amount of protein from each sample (5 μg) was dissolved in SDS-PAGE sample buffer (50 mM Tris-HCL pH 6.8, 2% SDS, 10% glycerol, 5% β-mercaptoethanol, 0.005% bromophenol blue), heated at 90 °C for 5 min, loaded into acrylamide gels in a random order with respect to genotype and sex (8% gel for analysis of synGAP, GluN2B, NLG-1 and 2; or 12% for TARPs), fractionated, and electrically transferred to PVDF membranes in 25 mM Tris, 150 mM glycine at 250V for 2-3 hours at 4°C. Completeness of transfer was checked by staining of gels with Gel Code Blue (Thermo Fisher Scientific) after transfer. Membranes were washed for 5 mins three times in 20 mM Tris, pH 7.6, 150 mM NaCl (TBS), blocked with Odyssey blocking buffer (LI-COR Biosciences, Lincoln NE) for 45 mins at RT, washed for 5 mins three times in TBS plus 0.1% tween (TBST) and then incubated in primary antibodies dissolved in TBST plus 5% BSA ON at 4°C. Each blot was incubated with a mixture of mouse anti-PSD-95 (50% ammonium sulfate cut of Ascites fluid of 7E3-1B8, AB_212825, dilution 1:10,000), and one of the following; rabbit anti-SynGAP (Pierce PA1-046, AB_2287112 dilution 1:3500), rabbit anti-TARP (EDM Millipore Ab9876, AB_877307 dilution 1:300), rabbit anti-GluN2B (raised in our lab, Zhou et al., 2007, see Fig. S1, dilution 1:1000), rabbit anti-NLG-1 (Synaptic Systems 129013 AB_2151646 dilution 1:2000), or rabbit anti-NLG-2 (Synaptic System 129202 dilution 1:1000). Membranes were washed for 5 min three times in TBST and incubated for 45 minutes at RT in 5% nonfat milk in TBST with a mixture of secondary antibodies including Alexa Fluor 680 goat anti-mouse IgG (Thermo Fisher Scientific A28183, AB_2536167, 1 μg/ml) to label PSD-95, and IRdye800-conjugated goat-anti-rabbit IgG (Rockland Immunochemicals, Limerick, PA, 611-145-122, AB_1057618, 1 μg/ml) to label the target proteins. Membranes were then given three 5 min washes in TBST, followed by three 5 min washes in TBS. Bound antibodies were visualized with the Odyssey Infrared Imaging System (LI-COR Biosciences).

### Data Quantification and Analysis

For each gel lane (representing a single animal) the ratio of the target protein (NLG-1, NLG-2, TARPs, GluN2B, or synGAP) to PSD-95 was calculated. Regions of Interest (ROI) were drawn around each protein band and the intensity within the box was determined with the use of Image Studio Light software supplied by LI-COR. To measure background, a box of the same size was placed in the lane in an unstained region above or below the band of interest. The intensity values were transferred to Microsoft Excel. For both the target protein and PSD-95, background was subtracted from the signal, then the ratio of the two background-corrected signals was calculated. At least three technical replicates were performed for each animal for each target.

To gather the data, cohorts of 7.5 week old mice, 7.5 week old rats, 12.5 week old mice, and 12.5 week old rats were processed separately. Because of the large number of animals within each cohort (∼40), gels were run and the samples were quantified over a few days and sometimes with different lots of antibodies. To normalize intensity signals within each cohort between different days and different lots of antibodies for Figs. 1 and 3, we first grouped the ratios for each target (e.g synGAP/PSD-95) according to the genotype and sex of the animals. For each combination of genotype and sex (e.g. WT male), we averaged the ratios determined for each target on each day, then calculated the overall average of these daily averages. To obtain correction factors that compensated for systematic small variations in signals on different days and for different lots of antibodies, we calculated the ratio of each daily average to the overall average.

Each ratio for individual animals (ie. from a single lane) was then adjusted by multiplying by the appropriate correction factor. Eighty percent of the correction factors fell between 0.5 and 1.75. One percent were larger than 2.75 and four percent were less than 0.5. After normalized within cohorts, the ratios were used for the analyses in figures 1 and 3. This procedure allowed us to correct for technical variation within cohorts while preserving true differences in ratios based on species, age, genotype, or sex. Analyses for significance were carried out and graphs were created with Prism 8 (GraphPad Software, San Diego). The D’Agostino-Pearson omnibus test was used to determine the normality of the data sets. The means of groups of data were compared for significant differences with either a one-tailed Mann-Whitney test (when non-normal), or a one-tailed paired T-Test, as indicated. If the variances of the two groups were found to be different using an F test, a one-tailed T-Test with a Welch’s correction was applied.

To test rigorously for correlations among individual animals between the ratio of synGAP to PSD-95 and the ratio of the other target proteins to PSD-95 (Figs. 2, 4, and 5), an additional normalization was applied with the use of Excel. We corrected for differences in intensity of signals between the four cohorts (i.e. 7.5 week old mice, 7.5 week old rats, 12.5 week old mice, and 12.5 week old rats) which had been analyzed separately. (The previous normalization only corrected for technical variation within each cohort.) Data for each cohort was divided into ratios from WT males, WT females, HET males, and HET females. Averages of the ratios for each protein within each of these groups, were calculated. The averaged ratios for each group were then further averaged across the cohorts. A normalization factor was calculated for each protein in each cohort by dividing the average for the cohort by the overall average for all the cohorts. Then the appropriate normalization factor was applied to individual data points in each cohort. This sequence corrected for variation in the average intensities of signals between cohorts and groups (for example, the overall lower expression of TARPS in 7.5 week old HET females [Fig. 1B]) and allowed us to look for correlations between ratios among the individuals across cohorts. To determine the correlations between ratios shown in Figs. 2, 4, and 5, we treated data as non-normal and calculated one-tailed Spearman rank correlation coefficients with Prism software.

## Acknowledgements

This work was supported by U. S. National Institute of Mental Health Grant MH115456 to MBK, the Allen and Lenabelle Davis Foundation (MBK), U.S. National Science Foundation Fellowship 1612289 to TLM, the Department of Biotechnology, India (SC and PCK), Simons Foundation Autism Research Initiative Grant 529085 to PK, Patrick Wild Centre (SMT and PCK), and Medical Reasearch Council UK Grant MR/P006213/1 to SMT and PCK.

## Supplementary Figures

**Fig. 2S1.**
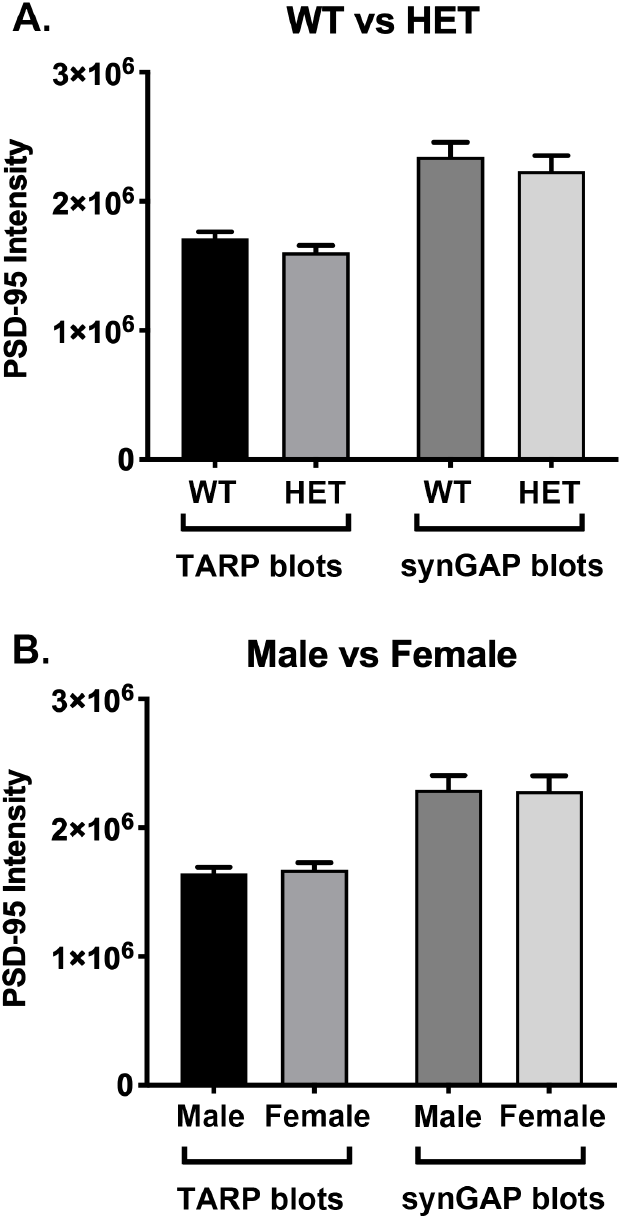
Intensity of PSD-95 bands on immunoblots. Blots were double-stained for PSD-95 and the target proteins as described under Methods. SynGAP, GluN2B, NLG-1 and NLG-2 were fractionated on 8% gels. TARPS were fractionated on 12% gels. The averaged intensity of staining of PSD-95 was not statistically different between WT and HET (A) or between Male and Female (B), measured on blots of 8% or 12% gels.

**Fig. 2S2.**
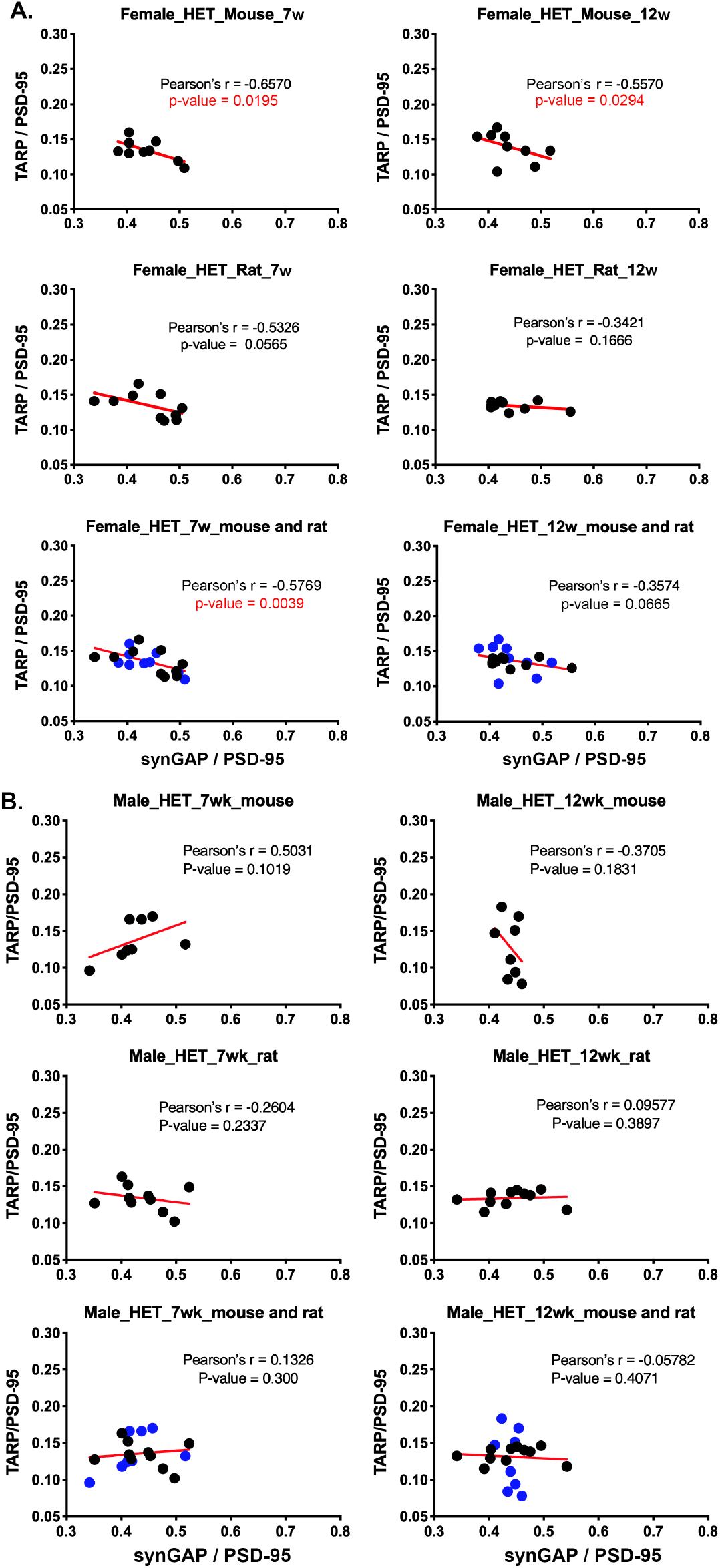
Correlation analysis between synGAP/PSD-95 and TARP/PSD-95 for data from 7 and 12 week old mice and rats. Data was acquired as described under Methods and in Fig. 2. The small individual data sets indicated in the headings of each panel were normal in distribution. Therefore, Pearson’s r, the parametric equivalent of the non-parametric Spearman’s coefficient, was calculated for each set. A) A significant inverse correlation was found for 7 and 12 week old female mice, and a strong trend was present for 7 and 12 week old female rats. B) None of the data sets containing only males showed a significant inverse correlation. P-values indicating statistical significance are shown in red.

## References

Berryer, M.H., Hamdan, F.F., Klitten, L.L., Moller, R.S., Carmant, L., Schwartzentruber, J., Patry, L., Dobrzeniecka, S., Rochefort, D., Neugnot-Cerioli, M., et al. (2013). Mutations in SYNGAP1 cause intellectual disability, autism, and a specific form of epilepsy by inducing haploinsufficiency. Hum Mutat 34, 385–394, doi: 10.1002/humu.22248.

Chen, H.-J., Rojas-Soto, M., Oguni, A., and Kennedy, M.B. (1998). A synaptic Ras-GTPase activating protein (p135 SynGAP) inhibited by CaM Kinase II. Neuron 20, 895–904

Cho, K.-O., Hunt, C.A., and Kennedy, M.B. (1992). The rat brain postsynaptic density fraction contains a homolog of the Drosophila discs-large tumor suppressor protein. Neuron 9, 929–942

Cohen, R.S., Blomberg, F., Berzins, K., and Siekevitz, P. (1977). The structure of postsynaptic densities isolated from dog cerebral cortex I. overall morphology and protein composition. J Cell Biol 74, 181–203

Dosemeci, A., Makusky, A.J., Jankowska-Stephens, E., Yang, X., Slotta, D.J., and Markey, S.P. (2007). Composition of the synaptic PSD-95 complex. Mol Cell Proteomics 6, 1749–1760, doi: 10.1074/mcp.M700040-MCP200.

Gray, N.W., Weimer, R.M., Bureau, I., and Svoboda, K. (2006). Rapid redistribution of synaptic PSD-95 in the neocortex in vivo. PLoS Biol 4, e370

Hamdan, F.F., Daoud, H., Piton, A., Gauthier, J., Dobrzeniecka, S., Krebs, M.O., Joober, R., Lacaille, J.C., Nadeau, A., Milunsky, J.M., et al. (2011). De novo SYNGAP1 mutations in nonsyndromic intellectual disability and autism. Biol Psychiatry 69, 898–901, doi: 10.1016/j.biopsych.2010.11.015.

Hamdan, F.F., Gauthier, J., Spiegelman, D., Noreau, A., Yang, Y., Pellerin, S., Dobrzeniecka, S., Cote, M., Perreau-Linck, E., Carmant, L., et al. (2009). Mutations in SYNGAP1 in autosomal nonsyndromic mental retardation. N Engl J Med 360, 599–605, doi: 10.1056/NEJMoa0805392.

Irie, M., Hata, Y., Takeuchi, M., Ichtchenko, A., Toyoda, A., Hirao, K., Takai, Y., Rosahl, T.W., and Sudhof, T.C. (1997). Binding of neuroligins to PSD-95. Science 277, 1511–1515

Jain, A., Huang, G.Z., and Woolley, C.S. (2019). Latent Sex Differences in Molecular Signaling That Underlies Excitatory Synaptic Potentiation in the Hippocampus. J Neurosci 39, 1552–1565, doi: 10.1523/JNEUROSCI.1897-18.2018.

Kennedy, M.B. (2013). Synaptic signaling in learning and memory. Cold Spring Harb Perspect Biol 8, a016824, doi: 10.1101/cshperspect.a016824.

Kennedy, M.B., Beale, H.C., Carlisle, H.J., and Washburn, L.R. (2005). Integration of biochemical signalling in spines. Nat Rev Neurosci 6, 423-434, 10.1038/nrn1685.

Kim, J.H., Liao, D., Lau, L.-F., and Huganir, R.L. (1998). SynGAP: a synaptic RasGAP that associates with the PSD-95/SAP90 protein family. Neuron 20, 683–691

Komiyama, N.H., Watabe, A.M., Carlisle, H.J., Porter, K., Charlesworth, P., Monti, J., Strathdee, D.J., O’Carroll, C.M., Martin, S.J., Morris, R.G., et al. (2002). SynGAP regulates ERK/MAPK signaling, synaptic plasticity, and learning in the complex with postsynaptic density 95 and NMDA receptor. J Neurosci 22, 9721–9732

Kornau, H.-C., Schenker, L.T., Kennedy, M.B., and Seeburg, P.H. (1995). Domain interaction between NMDA receptor subunits and the postsynaptic density protein PSD-95. Science 269, 1737–1740, doi: 10.1126/science.7569905.

Li, D., Qiu, Z., Shao, Y., Chen, Y., Guan, Y., Liu, M., Li, Y., Gao, N., Wang, L., Lu, X., et al. (2013). Heritable gene targeting in the mouse and rat using a CRISPR-Cas system. Nat Biotechnol 31, 681-683, 10.1038/nbt.2661.

McMahon, A.C., Barnett, M.W., O’Leary, T.S., Stoney, P.N., Collins, M.O., Papadia, S., Choudhary, J.S., Komiyama, N.H., Grant, S.G., Hardingham, G.E., et al. (2012). SynGAP isoforms exert opposing effects on synaptic strength. Nat Commun 3, 900, doi: 10.1038/ncomms1900.

Opazo, P., and Choquet, D. (2011). A three-step model for the synaptic recruitment of AMPA receptors. Mol Cell Neurosci 46, 1–8, doi: 10.1016/j.mcn.2010.08.014.

Sheng, M., and Kim, E. (2011). The postsynaptic organization of synapses. In Cold Spring Harb Perspect Biol, pp. a005678.

Till, S.M., Wijetunge, L.S., Seidel, V.G., Harlow, E., Wright, A.K., Bagni, C., Contractor, A., Gillingwater, T.H., and Kind, P.C. (2012). Altered maturation of the primary somatosensory cortex in a mouse model of fragile X syndrome. Hum Mol Genet 21, 2143-2156, 10.1093/hmg/dds030.

Tolias, K.F., Bikoff, J.B., Burette, A., Paradis, S., Harrar, D., Tavazoie, S., Weinberg, R.J., and Greenberg, M.E. (2005). The Rac1-GEF Tiam1 Couples the NMDA Receptor to the Activity-Dependent Development of Dendritic Arbors and Spines. Neuron 45, 525–538

Tomita, S., Chen, L., Kawasaki, Y., Petralia, R.S., Wenthold, R.J., Nicoll, R.A., and Bredt, D.S. (2003). Functional studies and distribution define a family of transmembrane AMPA receptor regulatory proteins. J Cell Biol 161, 805–816, doi: 10.1083/jcb.200212116.

Tomita, S., Stein, V., Stocker, T.J., Nicoll, R.A., and Bredt, D.S. (2005). Bidirectional synaptic plasticity regulated by phosphorylation of stargazin-like TARPs. Neuron 45, 269–277, doi: 10.1016/j.neuron.2005.01.009.

Vazquez, L.E., Chen, H.J., Sokolova, I., Knuesel, I., and Kennedy, M.B. (2004). SynGAP regulates spine formation. J Neurosci 24, 8862–8872, doi: 10.1523/JNEUROSCI.3213-04.2004.

Walkup, W.G., Mastro, T.L., Schenker, L.T., Vielmetter, J., Hu, R., Iancu, A., Reghunathan, M., Bannon, B.D., and Kennedy, M.B. (2016). A model for regulation by SynGAP-alpha1 of binding of synaptic proteins to PDZ-domain ‘Slots’ in the postsynaptic density. Elife 5, e16813, 10.7554/eLife.16813.

Walkup, W.G.t., Washburn, L., Sweredoski, M.J., Carlisle, H.J., Graham, R.L., Hess, S., and Kennedy, M.B. (2015). Phosphorylation of synaptic GTPase-activating protein (synGAP) by Ca^2+^/calmodulin-dependent protein kinase II (CaMKII) and cyclin-dependent kinase 5 (CDK5) alters the ratio of its GAP activity toward ras and rap GTPases. J Biol Chem 290, 4908–4927, doi: 10.1074/jbc.M114.614420.

Wang, W., Le, A.A., Hou, B., Lauterborn, J.C., Cox, C.D., Levin, E.R., Lynch, G., and Gall, C.M. (2018). Memory-Related Synaptic Plasticity Is Sexually Dimorphic in Rodent Hippocampus. J Neurosci 38, 7935–7951, doi: 10.1523/JNEUROSCI.0801-18.2018.

Zhou, Y., Takahashi, E., Li, W., Halt, A., Wiltgen, B., Ehninger, D., Li, G.D., Hell, J.W., Kennedy, M.B., and Silva, A.J. (2007). Interactions between the NR2B receptor and CaMKII modulate synaptic plasticity and spatial learning. J Neurosci 27, 13843–13853, doi: 10.1523/JNEUROSCI.4486-07.2007.

Zhu, J.J., Qin, Y., Zhao, M., Van Aelst, L., and Malinow, R. (2002). Ras and Rap control AMPA receptor trafficking during synaptic plasticity. Cell 110, 443–455, doi:10.1016/S0092-8674(02)00897-8.

